# Novel Compensatory Mechanisms Enable the Mutant KCNT1 Channels to Induce Seizures

**DOI:** 10.1101/191171

**Authors:** Salleh N. Ehaideb, Gentry T. Decker, Petrina Smith, Daniel Davis, Bing Zhang

## Abstract

Mutations in the sodium-activated potassium channel (KCNT1) gene are linked to epilepsy. Surprisingly, all KCNT1 mutations examined to date increase K^+^ current amplitude. These findings present a major neurophysiological paradox: how do gain-of-function KCNT1 mutations expected to silence neurons cause epilepsy? Here, we use *Drosophila* to show that expressing mutant KCNT1 in GABAergic neurons leads to seizures, consistent with the notion that silencing inhibitory neurons tips the balance towards hyperexcitation. Unexpectedly, mutant KCNT1 expressed in motoneurons also causes seizures. One striking observation is that mutant KCNT1 causes abnormally large and spontaneous EJPs (sEJPs). Our data suggest that these sEJPs result from local depolarization of synaptic terminals due to a reduction in Shaker channel levels and more active Na^+^ channels. Hence, we provide the first *in vivo* evidence that both disinhibition of inhibitory neurons and compensatory plasticity in motoneurons can account for the paradoxical effects of gain-of-function mutant KCNT1 in epilepsy.

## Introduction

Ion channels, such as potassium (K^+^), sodium (Na^+^), and calcium (Ca^2+^) channels, play important roles in neuronal excitability via regulation of the resting potential, the amplitude and the duration of action potentials (APs), the firing rates or patterns of APs, and synaptic transmission. Mutations in voltage-gated cation channels (Na^+^, Ca^2+^, and K^+^) that result in neuronal hyper-excitability are often linked to epilepsy^1-3^. One of the distinct exceptions is the sodium-activated potassium channel (KCNT1, also called *Slack or Slo2.2*), which is encoded by the KCNT1 gene and unique in that they are activated by intracellular Na^+^ and Cl^-4-8^. At least 24 point mutations have been found in the human *KCNT1* gene in patients with autosomal dominant nocturnal frontal lobe epilepsy (ADNFLE), malignant migrating partial seizures of infancy (MMIPSI), early onset epileptic encephalopathy (EOEE), and West syndrome^9-15^. The number of patients affected by KCNT1 mutations is not known but many of these patients do not respond well to current anti-epilepsy drugs^13,16-18^.

At present, the cellular mechanisms of KCNT1-related epilepsy remain poorly understood. Electrophysiological studies using heterologous expression systems such as the *Xenopus* oocytes or Chinese hamster ovary (CHO) cells have shown that all of the mutant KCNT1 channels studied to date significantly increase the magnitude of K^+^ currents^9,11,12,19^. These findings are important but they present an interesting neurophysiological conundrum in that increased K^+^ currents are expected to hyperpolarize or silence neurons and reduce the possibility for neurons to fire APs, a condition paradoxically unfavorable of triggering seizure or epilepsy.

Two different hypotheses have been proposed to explain this puzzle. The ‘repolarization hypothesis’^5,20-23^ states that increased Na^+^-dependent K^+^ currents might accelerate the rate of AP repolarization, thus enhancing the firing rate. Studies of KCNT1 KO mice showed, however, that the lack of the KCNT1 K^+^ current sped up repolarization and enhanced AP firing^24^. This result is not in agreement with the ‘repolarization hypothesis’, suggesting that KCNT1 K^+^ current hinders repetitive firing normally. Another hypothesis is the ‘disinhibition hypothesis’^19^, which states that increased K^+^ currents in inhibitory interneurons reduce GABA release and consequently cause hyperexcitability of postsynaptic neurons. This hypothesis is plausible; however, it has not been tested directly.

To test the disinhibition hypothesis and to discover new *in vivo* function of mutant KCNT1, we generated a fruit fly model of human mutant KCNT1. Our data show that expression of the mutant KCNT1 channels in GABAergic interneurons indeed causes bang-sensitive seizures in flies, the first *in vivo* observation in support of the disinhibition hypothesis. To our surprise, however, expression of the mutant channels in motoneurons results in uncoordinated larval locomotion, enhanced synaptic transmission, and also bang-sensitive seizures in adult flies. These results suggest that neuronal excitability is enhanced rather than reduced in motoneurons by the mutant KCNT1. We further show that neuronal hyperexcitability is achieved in part by a compensatory downregulation of endogenous *Drosophila* K^+^ channels such as Shaker and enhancement of voltage-gated Na^+^ channels, in an attempt to counter balance the effect of increased KCNT1 K^+^ current. Hence, our study supports two complementary hypotheses, the ‘disinhibition hypothesis’ and a new ‘compensatory plasticity hypothesis’, to help better understand the neurobiology underlying KCNT1-associated epilepsy.

## Results

### Mutant hKCNT1 channels increase K^+^ conductance in muscles

To date, nearly all electrophysiological studies of mutant KCNT1 channels associated with epilepsy have been conducted in heterologous expression systems, such as the Xenopus oocytes or CHO cells^9,11,12,19^. These studies show that mutant hKCNT1 channels display a significant increase in K^+^ current magnitude compared to control^9,11,12,19^. To determine whether hKCNT1 mutations also exhibit gain-of-function (an increase in K^+^ conductance) phenotypes *in vivo*, we used the *GAL4/UAS* system to overexpress wild type (WT) human KCNT1 (hKCNT1) and two mutants, G288S (a point mutation in S5; next to the channel pore) and R928C (a point mutation near the NAD^+^ domain) (Suppl Fig. 1a). Our data show that the resting potential is significantly hyperpolarized and the input resistance dramatically reduced in muscles expressing either G288S or R928C *hKCNT1* mutations by *Mhc-GAL4* (Suppl Fig. 1b-e). The *hKCNT1* mutant larvae were not able to crawl very far from its starting point and for the most part of the recording period the larvae were stationary and inactive (Suppl Movie 1 and Suppl Fig. 2). These results are similar to the effects of human inward-rectifier potassium channel 2.1 (*Kir2.1*)^25^ expressed in muscles (Suppl Fig. 1b-e). Thus, our *in vivo* data illustrate that *hKCNT1* G288S and R928C mutations are gain-of-function resulting in an increase in K^+^ conductance, and subsequently leading to impaired larvae locomotion activities, hyperpolarized resting potential, and lower muscle resistance.

### Mutant KCNT1 causes seizures in adult flies

Epilepsy is neurological disease defined as involuntary muscle convulsions resulting in seizures. Drosophilists have been using adult flies to determine the genetic bases of epilepsy and whether a particular mutation will predispose the flies to seizures. Several behavior and electrophysiological techniques have been developed to assess seizures in adult flies^26-28^. The bang-sensitivity behavioral assay is used to determine whether subjecting flies to mechanical stimulation can elicit seizure activities^29,30^.

To directly test the disinhibition hypothesis, we expressed the mutant and wildtype human KCNT1 in GABAergic neurons using a cell-specific *Gal4*, *Gad-Gal4*^31^. Adult flies were then subjected to the bang-sensitive test in which the flies were placed in an empty fly vial, vortexed for 20 sec, and then observed for seizing behavior (often flip on their backs and unable to stand up or buzz around for at least one second). We noted that the flies expressing mutant R928C in GABAergic neurons had a higher propensity to display seizure (Fig. 1a-e; Suppl Movie 2). The frequency of seizures is significantly higher and duration significantly longer compared to the two control groups (*Gad-Gal4 >WT*; *Gad-Gal4 > WT human KCNT1*). The mean seizing frequency is 26% compared to the controls (10%, 5%, respectively). The mean seizing duration of these flies is significantly longer (p<0.01), averaging 28 Sec compared to 2 sec and 1.5 sec in *Gad-Gal4>WT* and *Gad-Gal4>WT human KCNT1*, respectively. These results support the ‘disinhibition hypothesis’, indicating that silencing inhibitory neurons tips the balance towards excitation and thus implicating a role for GABAergic neurons in epilepitogenesis in human patients.

We were also curious about whether expression of the mutant KCNT1 channels in motoneurons would have a similar effect as they did in GABAergic neurons. In motoneurons mutant KCNT1 driven by *D42-Gal4*^32,33^ causes embryonic lethality at 25°C, consistent with the silencing effect of enhanced K^+^ current and pan neuronal expression of D42 in early embryos. To overcome this obstacle, we used *GAL80*^*ts*^ ^34^ (this form of *GAL80* become inactive when flies are reared at higher temperature) in the context of the *D42, UAS hKCNT1* mutation to prevent the expression of hKCNT1 channels during development. Flies were kept at 18 °C until eclosion, then adult flies were collected and kept at 25 °C to age, allowing the expression of hKCNT1 mutant channels in adult flies. Mechanical stimulation of flies expressing R928C mutation by vortexing showed a significant increase in the percentage of seizing flies and seizing duration (25.7 ± 3.5 S and 11.3 ± 2.1 S, respectively) compared to *D42/+* (1.25 ± 0.9 S and 1 S, respectively) and *D42/hKCNT1* (0.7 ± 0.7 S and 1 S, respectively, Fig. 1f-j and Suppl Movie 3). These data suggest a new mode by which mutant hKCNT1 channels cause seizure and epilepsy.

**Figure 1:**
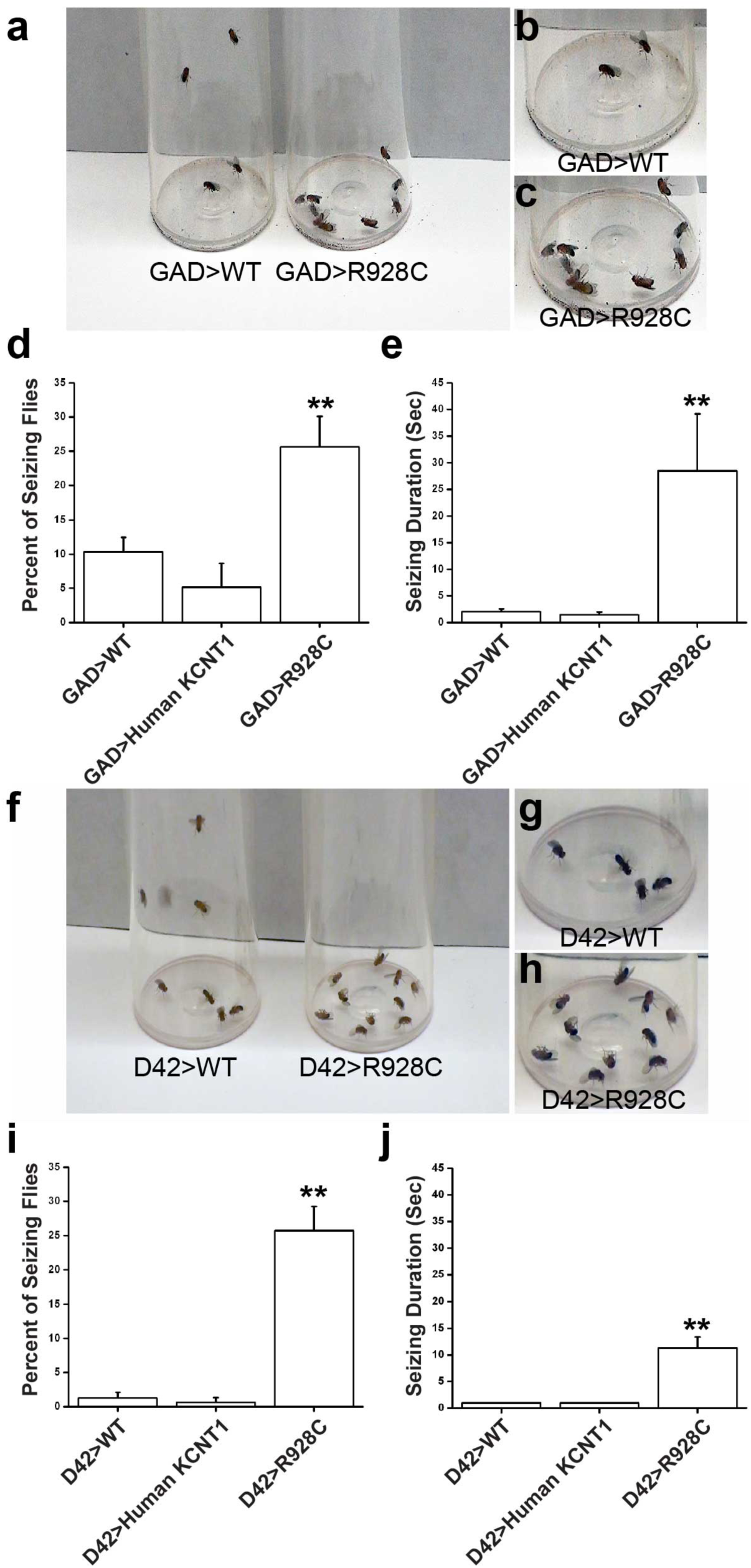
Neuronal expression of mutant hKCNT1 channel leads to seizures in adult flies. (**a, f**) Representative image of flies after twenty seconds of mechanical stimulation (vortexing). Close-up images of the bottom of the vials are shown in (**b, c, g**, and **h**). Control flies show no sign of seizure activity after vortexing whereas a large number of mutant KCNT1-expressing flies exhibit seizure. (**d** and **e**) GABAergic expression of mutant hKCNT1 channel in adult flies show a significant increase in both number of seizing flies and seizing duration. (**i** and **j**) Motoneuron expression of mutant hKCNT1 channel predispose flies to seizure and very few control flies seized after vortex. If control flies seized they did not seize more than one second while flies expressing mutant hKCNT1 channels showed significantly longer seizure durations. At least 120 and 40 flies were tested for Motoneuron and GABAergic expression per genotype. Error bars show S.E.M. \*\**P* < 0.01 one-way ANOVA, Turkey HSD *post hoc test*.

### Mutant hKCNT1 channels do not silence motoneurons but disrupt coordinated crawling and motoneuronal firing

The findings from expressing mutant KCNT1 in muscle, GABAergic neurons, and motoneurons appear to contradict one another in that muscles and GABAergic neurons are silenced whereas motoneurons in adult flies are not. In order to better understand the mechanisms by which mutant KCNT1 in motoneurons causes seizures, we decided to examine the impact of mutant KCNT1 on larval behavior and physiology. To overcome the embryonic lethality, we raised the flies with motoneuronal expression of mutant KCNT1 at 18°C, which reduces the expression levels of the mutant hKCNT1 channels. *D42>G288S* reached pupal stage and few of the *D42>R928C* flies were able to reach the adult stage. We noted that larvae expressing the mutant KCNT1 channels showed severe crawling defects compared to control larvae (wildtype or larvae expressing the wildtype KCNT1) (Suppl Fig. 3a,b; Suppl Movie 4). Because of the uncoordinated crawling, the pupae became bended (crescent shape, Suppl Fig. 3c). However, after careful observation we noted that unlike muscle expression of mutant hKCNT1 channels, which were inactive for the most part of the observation, larvae expressing G288S or R928C in motor neurons showed a strong and active but one sided larval contractions (Suppl Movie 4). This finding suggests that the reduction in larvae crawling distance is not due to neuronal inactivity but rather uncoordinated neuronal firing.

Our behavioral observations of both larvae and adults strongly suggest that neuronal expression of mutant KCNT1 at low levels does not silence motoneurons. On the contrary, they cause hyperactivity in larvae (see below) and seizures in adult. To determine the effects of *hKCNT1* mutations on neuronal activity, we utilized the well-established neuromuscular junction (NMJ) preparation to assess their effects on synaptic transmission in third-instar larvae^35,36^. Surprisingly, electrical stimulation of the segmental nerve did not alter the amplitude of excitatory junction potentials (EJPs) of larvae expressing G288S or R928C mutations in motor neurons compared to control larvae. The mean EJP amplitude for *D42-GAL4>G288S*, *D42>R928C* was 49.1 ± 1.7 mV and 49.5 ± 1.7 mV, respectively, which were not significantly different from *D42/+* (50.6 ± 1.1 mV) and *D42/hKCNT1* (49.5 ± 1.3 mV) (Fig. 2a,b). In addition, there was no significant difference in membrane resting potentials between of G288S and R928C mutations and controls (*D42/+* and *D42/hKCNT1*) (Suppl Fig. 4). These data indicate that reduction of larval crawling activity caused by motoneuronal expression of mutant hKCNT1 channels is not due to neuronal silencing but most likely due to inability to coordinate neuronal firing, which hinders larvae crawling ability.

To more directly investigate whether *hKCNT1* mutations compromise coordinated neuronal firing, we examined ventral nerve cord (VNC) neuronal firing in third-instar larvae using Ca^2+^ imaging. Larvae VNC (equivalent to spinal cord in humans) have eight pairs of abdominal ganglia, each right and left ganglion controls the corresponding larval body wall segment^37-40^. For example, the first pair of ganglia control the right and left abdominal muscle segment 1. During larvae crawling the first pair of ganglia will fire together then the next pair and signal will propagate down until the last pair fired^37-40^. To visualize ganglia firing we used the calcium sensor *GCaMP6S* expressed in motoneurons. Larvae expressing WT hKCNT1 channels showed coordinated ganglion firing, where right and left ganglia fired simultaneously (Fig 2c,d and Suppl Movie 5). On the other hand, expression of hKCNT1 mutated channels resulted in a significant number of unsynchronized motoneuronal firing, and sometimes only ganglion in one side fired without the other pair firing at all (Fig. 2c,d,e and Suppl Movie 5). In addition, oftentimes calcium waves were not able to pass through all ganglia and usually terminated after reaching the fourth ganglion in larvae expressing mutant hKCNT1 channels. However, both larvae expressing WT and mutated hKCNT1 channels showed a similar peak amplitude of Ca^2+^ waves (Fig. 2f). Taken together, our data on both EJP and calcium waves suggest that hKCNT1 mutations do not silence neuronal firing but rather influence firing synchrony.

**Figure 2:**
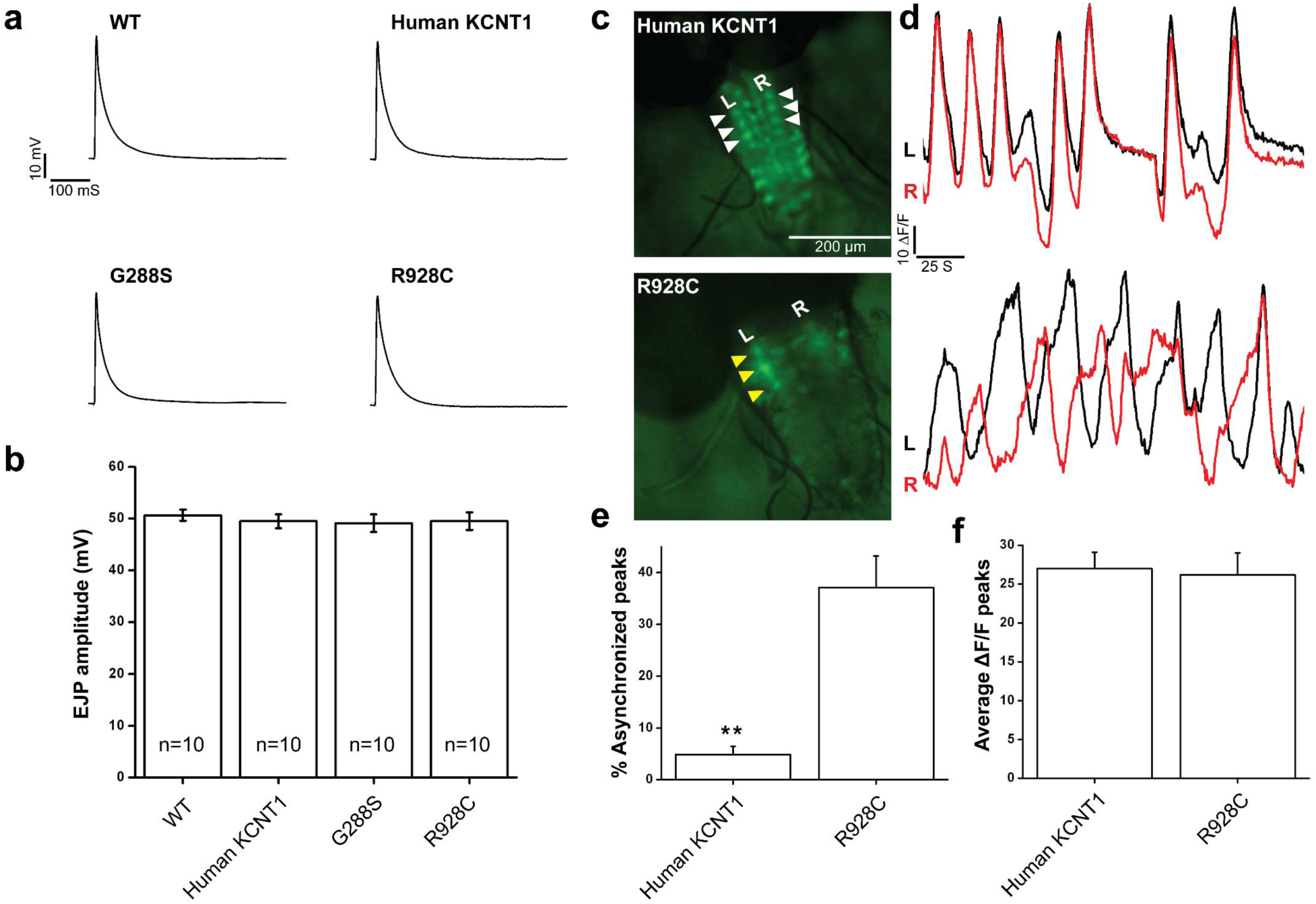
Expression of *hKCNT1* mutation did not silence neurons, but caused uncoordinated ventral ganglion firing. (**a**) Representative traces of excitatory junction potential (EJPs) under physiological conditions. (**b**) Average EJPs amplitude of ten recordings (two per larva) per genotype. Recordings were taken from muscle 6 segment A3 or A4. (**c**) Representative image of calcium waves in larvae expressing WT hKCNT1 channels (top) in motor neurons and larvae expressing mutated hKCNT1 channel (bottom). The calcium waves were visualized using *GCaMP6S* calcium sensor expressed in motoneuron. **L** and **R** indicate left and right sides, respectively, of ventral nerve cord (VNC) in third-instar larvae. White and yellow arrow heads indicate coordinated and uncoordinated left and right VNC ganglion firing respectively. (**d**) Representative calcium waves signals from left and right VNC ganglion pairs, showing coordinated (top) and uncoordinated (bottom) waves in larvae expressing WT and mutated hKCNT1 channels during active crawling respectively. (**e**) Quantification of percent of uncoordinated left and right VNC ganglion calcium waves. (**f**) Quantification of average calcium wave amplitude. Five larvae were used for calcium waves analysis. All larvae were reared at 22 °C for 5 days and 2 days at 25°C prior to EJP recordings and calcium imaging. Error bars show S.E.M. \*\**P* < 0.01, one-way ANOVA, *Student’s T test*.

### Mutant hKCNT1 channels cause large spontaneous postsynaptic potentials at the larval NMJ

Spontaneous miniature excitatory junction potentials (mEJPs or minis) play a role in synaptic plasticity and function, evoked transmitter release, neuronal excitability, and postsynaptic membrane resistance^41-45^. It has been shown that in somatostatin cells taken from epileptic mice exhibit an increase in miniature excitatory postsynaptic synaptic current (mEPSC) frequency^46^. Thus, we investigated whether *hKCNT1* mutations alter the mini properties at the NMJ of third-instar larvae. We severed the segmental nerves posterior to the ventral ganglion and monitored mEJP activity (usually spontaneous release from single vesicles) at the NMJ.

Control larvae (both *D42/+* and *D42/hKCNT1*) displayed typical spontaneous minis, with an average frequency of 1.4 Hz and amplitude of 1.7 ± 0.1 mV (for both genotypes). Remarkably, larvae expressing mutant hKCNT1 channels showed very large spontaneous synaptic potentials, up to 14 mV (Fig. 3a). As shown below, these large synaptic events are spontaneous EJPs. Hence, we will refer them as spontaneous EJPs (sEJPs) rather than minis or mEJPs. The average amplitude of spontaneous synaptic potentials (note that this refers to both minis and sEJPs) in larvae expressing G288S and R928C mutations was significantly higher (2.6 ± 0.2 mV and 3 ± 0.2 mV, respectively, ANOVA, *P* < 0.01). The counts of minis plus sEJPs and their cumulative probability plots are shifted to the right in larvae expressing mutant hKCNT1 channels, due to the presence of significantly high number of large sEJPs in these larvae compared to *D42/+* and *D42/hKCNT1* larvae (Fig. 3b,c).

**Figure 3:**
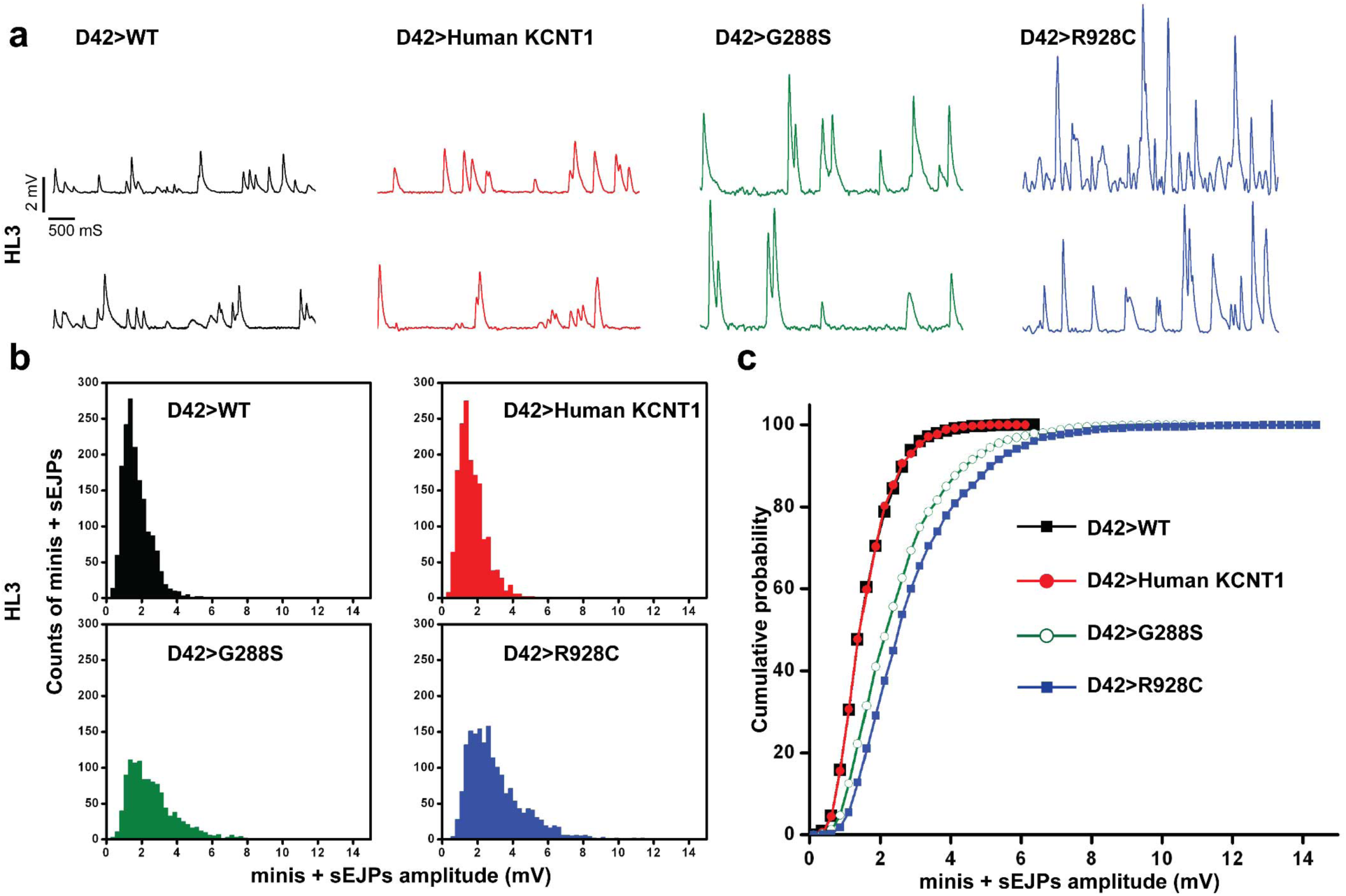
Third-instar larvae expressing *hKCNT1* mutations in motor neurons showed spontaneous EJPs. (**a**) Representative traces of miniature excitatory junction potentials (mEJP or minis) under physiological condition. Both *hKCNT1* mutants show significantly unusually larger spontaneous synaptic events in addition to minis compare to *D42/+* and *D42/hKCNT1* traces. We call these large synaptic potentials spontaneous EJPs (sEJPs). (**b**) Histograms of spontaneous synaptic potentials (minis plus sEJPs) of the four different genotypes. Synaptic potential counts are sorted into 0.125 mV bins. Note the right-shift distribution of spontaneous synaptic potentials in the two mutant KCNT1 larvae. (**c**) Cumulative probability plot of spontaneous synaptic potentials from larvae expressing mutant hKCNT1 channels illustrate the present of significantly higher number of large sEJPs compare to *D42/+* and *D42/hKCNT1*. Recordings were taken from muscle 6 segment A3 or A4 and all larvae were reared at 22 °C for 5 days and 2 days at 25 °C prior to synaptic recordings. A total of ten recordings (two per larva) were made per genotype.

These sEJPs are highly unusual as the axons of the motoneurons are cut free from the soma and it is not possible for action potentials to be propagated from the soma. We reasoned that they were caused by local depolarization at the synaptic terminal. This could be possible only if there was a compensatory enhancement in voltage-gated cation (Na^+^ or Ca^2+^) channel levels and activities or a reduction in K^+^ channel levels and activities. To test this hypothesis, we used tetrodotoxin (TTX) to block voltage-gated sodium channel activity and surprisingly observed that TTX only reduced the number of ‘extremely’ large sEJPs (>10 mV) but did not abolish sEJPs in larvae expressing mutant hKCNT1 channels (Fig. 4a,b,c; Suppl Fig. 5). However, we noticed a slight decrease in average amplitude of spontaneous synaptic potentials in the G288S and R928C expressing larvae (2.1 ± 0.1 mV and 2.5 ± 0.2 mV, respectively, ANOVA, *P* < 0.01), but TTX had no significant effect on mini amplitude in *D42/+* and *D42/hKCNT1* larvae (1.6 ± 0.1 mV and 1.5 ± 0.1 mV, respectively). These data suggest that there is an upregulation of voltage-gated Na^+^ channels but they are not responsible for eliciting the sEJPs.

We then added the Ca^2+^ channel blocker cadmium to the TTX-containing saline and effectively eliminated the sEJP in the mutant KCNT1-expressing larvae (Fig. 4d,e,f). On the other hand, no changes in mEJP properties were observed in the *D42/+* and *D42/hKCNT1* larvae. These results indicate that depolarization of voltage-gated Ca^2+^ channels at the NMJ likely results in more transmitter release and gives rise to sEJPs.

**Figure 4:**
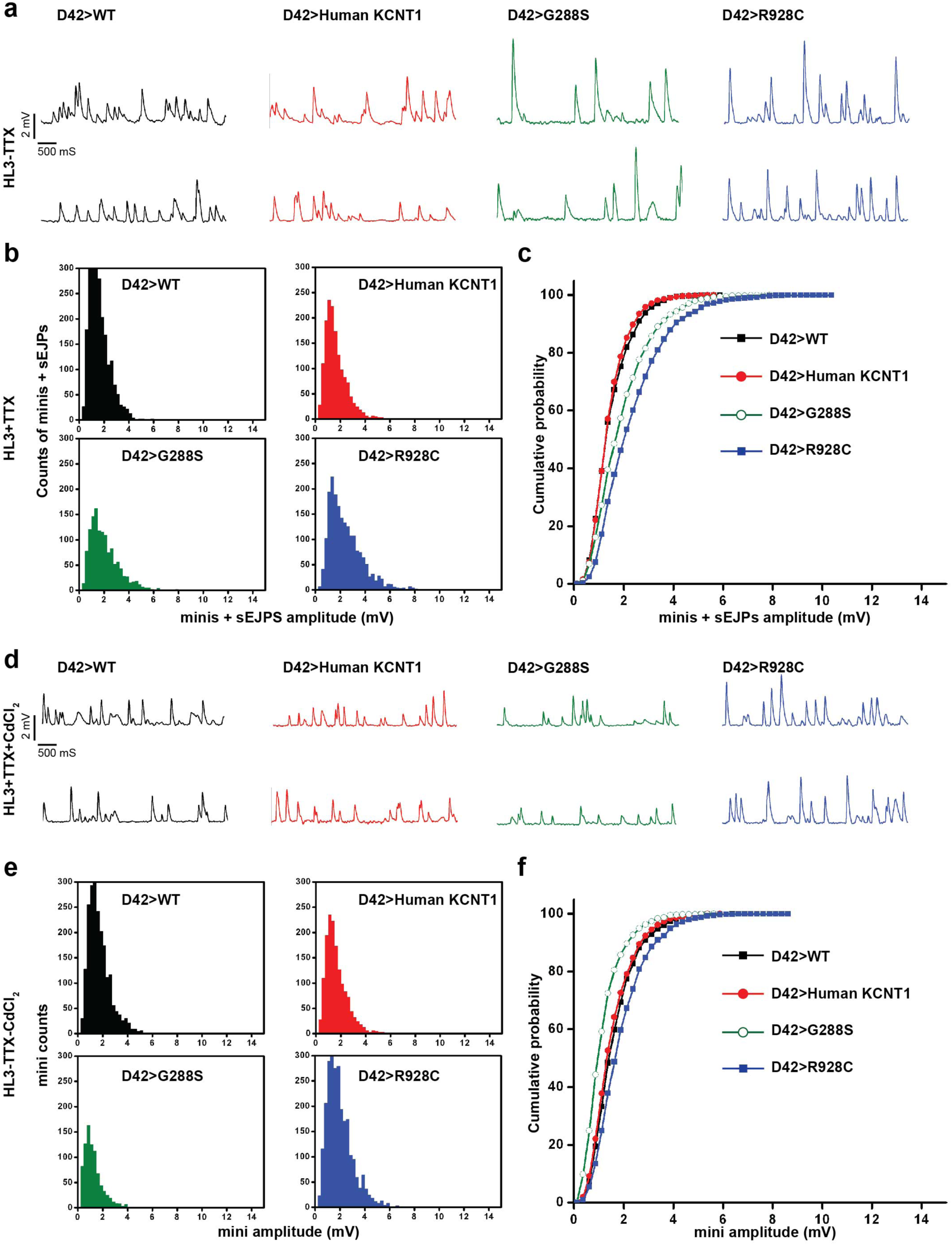
Blocking both voltage-gated sodium and calcium channels in larvae expressing mutant hKCNT1 channels eliminated the spontaneous EJPs. (**a**) Representative traces of spontaneous synaptic potentials (minis + sEJPs) treated with voltage-gated sodium channel blocker TTX (1 µM) in HL-3 saline. Note that the large sEJPs persist in TTX-saline. (**b** and **c**) Histograms and cumulative probability plots show that the extremely large sEJPS (>10 mV) are blocked by TTX but TTX does not eliminate all sEJPs. (**d**) Representative traces of spontaneous synaptic potentials (minis + sEJPs) treated with both voltage-gated Na^+^ channel blocker TTX (1 µM) and voltage-gated K^+^ channel blocker CdCl_2_ (10 µM) in HL-3 saline (This treatment successfully reduces or abolishes the occurrence of spontaneous EJPs. (**e** and **f**) Histograms and cumulative probability plots show similar distributions of spontaneous synaptic potentials in TTX and Cd^2+^-treated larval NMJs. Recordings were taken from muscle 6 segment A3 or A4 and all larvae were reared at 22 °C for 5 days and 2 days at 25 °C prior to mini recordings. A total of ten recordings (two per larva) were made per genotype.

### Immunocytochemistry reveals compensatory decreases in endogenous K^+^ channels in larvae expressing mutant KCNT1

Our behavioral, imaging, and electrophysiological studies collectively suggest that mutant KCNT1 channels trigger compensatory mechanisms by which other ion channels are down or up regulated to counter balance the silencing effect of KCNT1 K^+^ channels. In other words, the large increase in K^+^ currents through mutant KCNT1 channels has the potential to fully or partially silence neurons, like Kir2.1 does, to reduce the possibility of firing and impair synaptic transmission. In contrast, the motoneurons appear to fire normally (albeit not synchronized) and the EJP amplitude is largely unchanged. What enables the motoneurons to fire action potentials and the synaptic terminal to produce local and spontaneous auto-depolarization? We hypothesize that there is a compensatory change of excitability at both cell bodies and axons to ensure that action potentials can be produced and propagated. Furthermore, we hypothesize that there is a local compensation at the NMJ to enhance synaptic activity. We used antibodies specific to Shaker (Sh) to stain the CNS and the neuromuscular preparation of larvae and showed that Shaker channels are reduced in levels (Fig. 5a,b).

**Figure 5:**
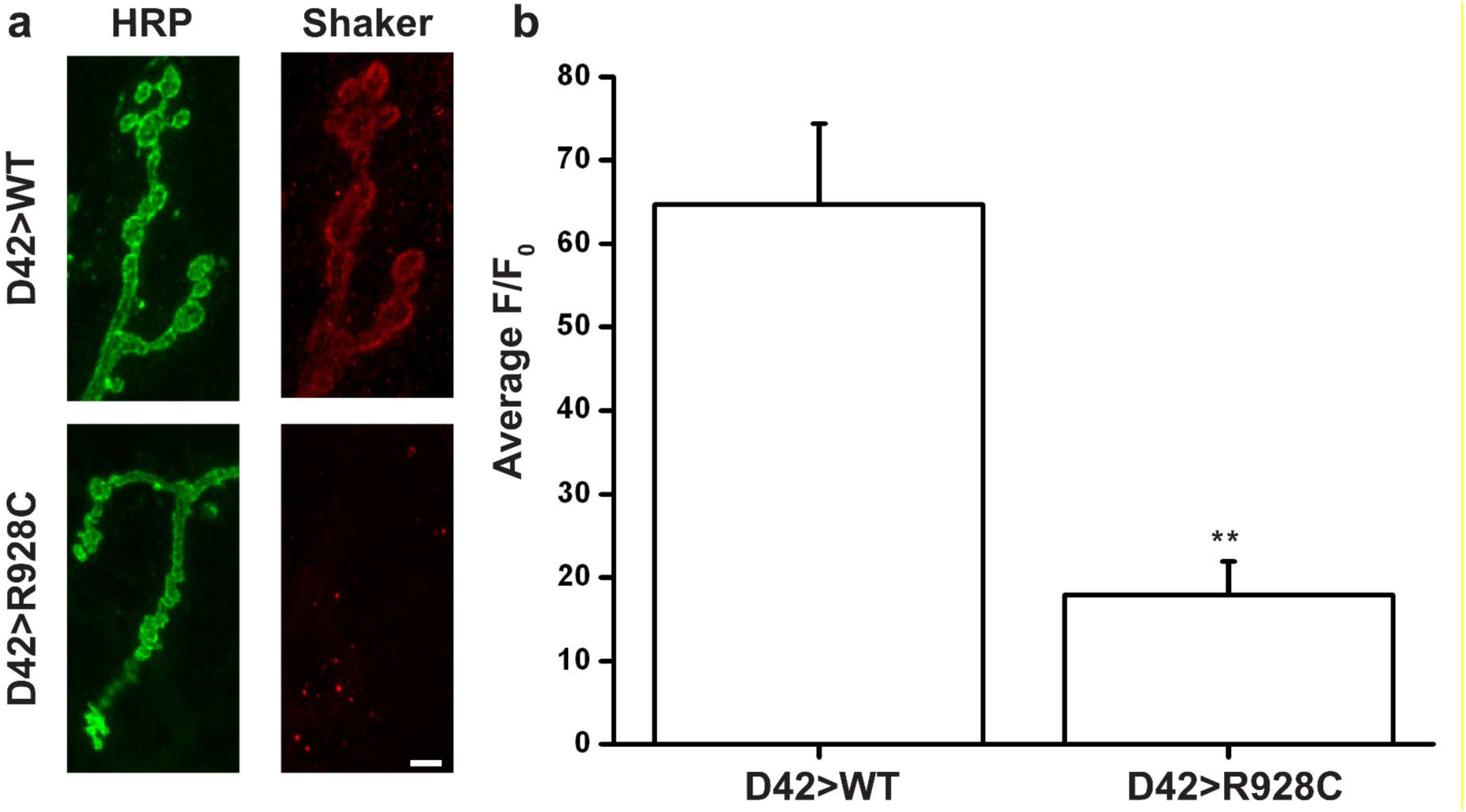
Neuronal expression of mutant hKCNT1 channel leads to significant reduction in Shaker channels staining at the NMJ. (**a**) Representative NMJ confocal images of WT and larvae expressing mutant hKCNT1 channel in motoneurons. Larvae expressing mutant hKCNT1 channel show weak or no Shaker staining. Scale = 5 µM. (**b**) Quantification of Shaker staining intensity show that Larvae expressing mutant hKCNT1 channel have a significant reduction in Shaker signal compared to control. \*\**P* < 0.01, *Student’s T test*.

## Discussion

The discovery of the link between mutant KCNT1 and epilepsy is important but it also presents challenges understanding the role of the K^+^ channel in epileptogenesis. *In vitro* studies to date all show that mutant KCNT1 channels significantly increases K^+^ current magnitude^9,11,12,19^. It is easier to understand why mutations enhancing Na^+^ and Ca^2+^ channel activities or reducing K^+^ channel activities can cause hyperexcitability and increase the probability for seizures or epilepsy. Similarly, mutant Cl^-^ channels in skeletal muscles both in the fainting goat and humans cause myotonia^47-49^. Neurophysiologically speaking, enhancing K^+^ currents is expected to hyperpolarize the resting potential or truncate action potentials and thereby reducing the possibility for neuronal firing. What then accounts for the neuropathology in KCNT1-associated epilepsy?

Two different hypotheses have been proposed to explain why enhanced K^+^ channel activities in mutant KCNT1 could cause hyperexcitability and generate conditions in favor of seizures. One hypothesis, which we name ‘repolarization hypothesis’, states that following Na^+^ influx the activation of KCNT1 K^+^ channels shortens the duration of APs by repolarizing it at a faster rate. This in turn enables the neuron to fire more APs per unit time, resulting in hyperexcitability. This possibility is plausible, as shown in electrocytes of some electrical fish^20^ and BK channel and β subunit -linked epilepsy^50-52^. However, it will depend on the mode by which Na^+^ activates KCNT1 channels and the kinetics of the K_(Na)_ current^53^. KCNT1 channels are unique in that is activated by intracellular Na^+^ and Cl^-^. Salkof and colleagues showed that activation of KCNT1 channels does not need high intracellular [Na^+^]_i_^54^, implying that KCNT1 may also be important for contributing to resting potential as well as repolarizing AP. Furthermore, K_(Na)_ following Na^+^ influx could outlast the duration of an AP, especially during afterhyperpolarization period, and therefore prevent or delay the onset of the next AP. Finally, genetic studies of KCNT1 KO in mice showed that AP repolarization is faster and neurons fire more APs^24^. This *in vivo* study indicates that the presence of KCNT1 K current normally hinders excitability or silence neurons. Our data obtained from muscle expression of the mutant KCNT1 provide strong evidence that the mutant KCNT1 channels are similar to inward rectifier K^+^ channel 2.1 in reducing the muscle input resistance and hyperpolarizing the muscle resting potential. Further, neuronal expression of the mutant KCNT1 at high levels causes embryonic lethality or folded wings (Suppl Fig. 6) if the flies live to adult. These observations are consistent with the notion that mutant KCNT1 are gain of function mutations that silence both muscles and neurons. Hence, studies in both mice and fruit flies do not support the ‘repolarization hypothesis’.

The second hypothesis, which we call the ‘disinhibition hypothesis’^19^, postulates that the enhanced mutant KCNT1 K^+^ current silences inhibitory neurons and thereby removing inhibition of neural circuits and tipping the balance towards hyperexcitability and seizures. This is an exciting hypothesis, but up to this point this hypothesis has not been directly tested. Our data showing that expression of mutant KCNT1 in GABAergic neurons induces seizures in adult flies lends a strong support to this hypothesis.

However, the disinhibition hypothesis alone may not be sufficient to account for the mutant KCNT1 actions in the nervous system. This is because KCNT1 is broadly expressed in a variety of neuronal types and regions in the human brain^55,56^. The broad and complex expression pattern of KCNT1 begs for additional mechanisms other than the disinhibition hypothesis to account for the effect of mutant KCNT1 in other neurons. By expressing the mutant KCNT1 in motoneurons we have learned that these neurons can be silenced if the expression levels are high or become hyperexcited if the expression levels are low. When specifically expressed in adult motoneurons, mutant KCNT1 also causes bang-induced seizures, suggesting that these neurons are not silenced. This is consistent with the observations of normal EJPs and Ca^2+^ peaks in motoneurons in larvae. More strikingly, we reveal novel changes at the synapse where local depolarization leads to spontaneous synaptic potentials, which is partially sensitive to TTX blockade and fully sensitive to Cd^+^. At the NMJ, the major endogenous K^+^ channel Shaker is significant reduced in levels, and TTX can block some of the sEJPs, providing a molecular explanation for the enhanced excitability in motoneurons. Our finding of sEJPs is consistent with previous findings that sEJPs also occurred in flies with mutations in Sh eag K^+^ channels^57^. Based on these observations, we propose a ‘compensatory plasticity hypothesis’ as a novel mechanism to counter balance the silencing effect of mutant KCNT1 currents as an additional means to produce neuronal hyperexcitability and seizures. This work adds to the growing list of examples of homeostatic regulation of neuronal excitability^58^, although in our KCNT1 fly model it fails to fully compensate to restore normal function. The disinhibition and compensatory mechanisms are complementary to each other, suggesting that mutant KCNT1 channels likely have different effects on different neurons. Further studies are expected to pinpoint the regulatory mechanisms by which mutant KCNT1 regulates Sh, Na^+^ and perhaps ion channel gene expression or trafficking.

## Supplementary Figure legends

**Supplementary Figure 1:**
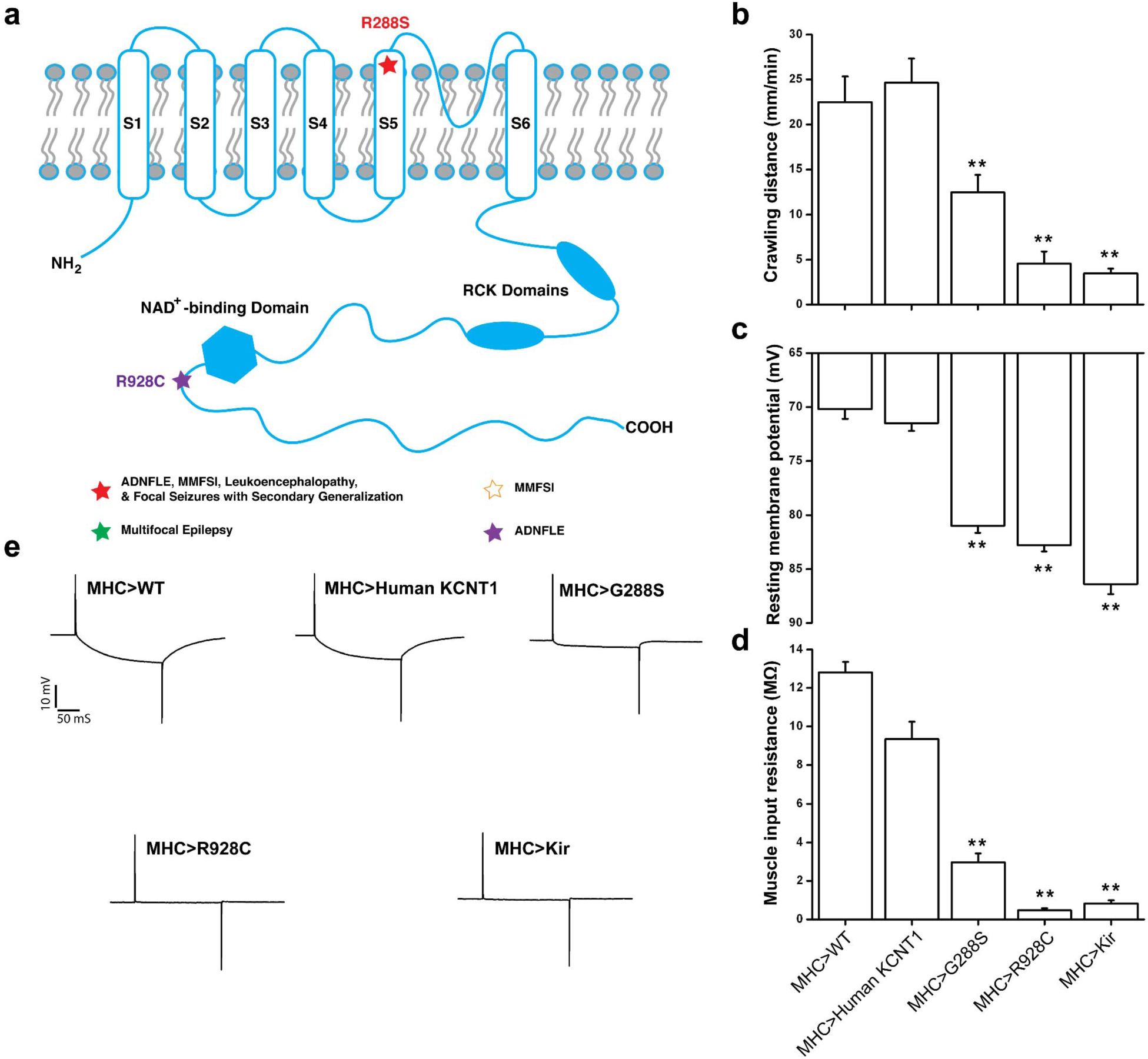
Muscle expression of mutant hKCNT1 protein in third-instar larvae reduce crawling distance, membrane resting potential, and muscle resistance. (**a**) Schematic diagram of human KCNT1 protein showing protein structure, location of the mutations studied in this paper, and type of epilepsy syndrome caused by these mutations. (**b**) Third-instar larvae expressing either mutant *hKCNT1* or *Kir2.1* in muscles show a significant reduction in average crawling distance compare to controls. At least twenty larvae were used per genotype (10 males and 10 females). (**c** and **d**) In addition, overexpression of mutant KCNT1 channels significantly hyperpolarized the average resting membrane potential (**c**) and average muscle input resistance (**d**). (**e**) Representative traces for muscle input resistance in control and experimental larvae. Note the dramatic and similar reduction in muscle input resistance in both *MHC>R982C* and *MHC>Kir2.1* larvae. Muscle 6 of abdominal segment 3 or 4 was used for assisting the resting membrane potential and muscle input resistance. Ten larvae were used per genotype. All larvae were kept at 25 °C until the time of testing. Error bars show S.E.M. \*\**P* < 0.01, one-way ANOVA, Turkey HSD *post hoc test*.

**Supplementary Figure 2:**
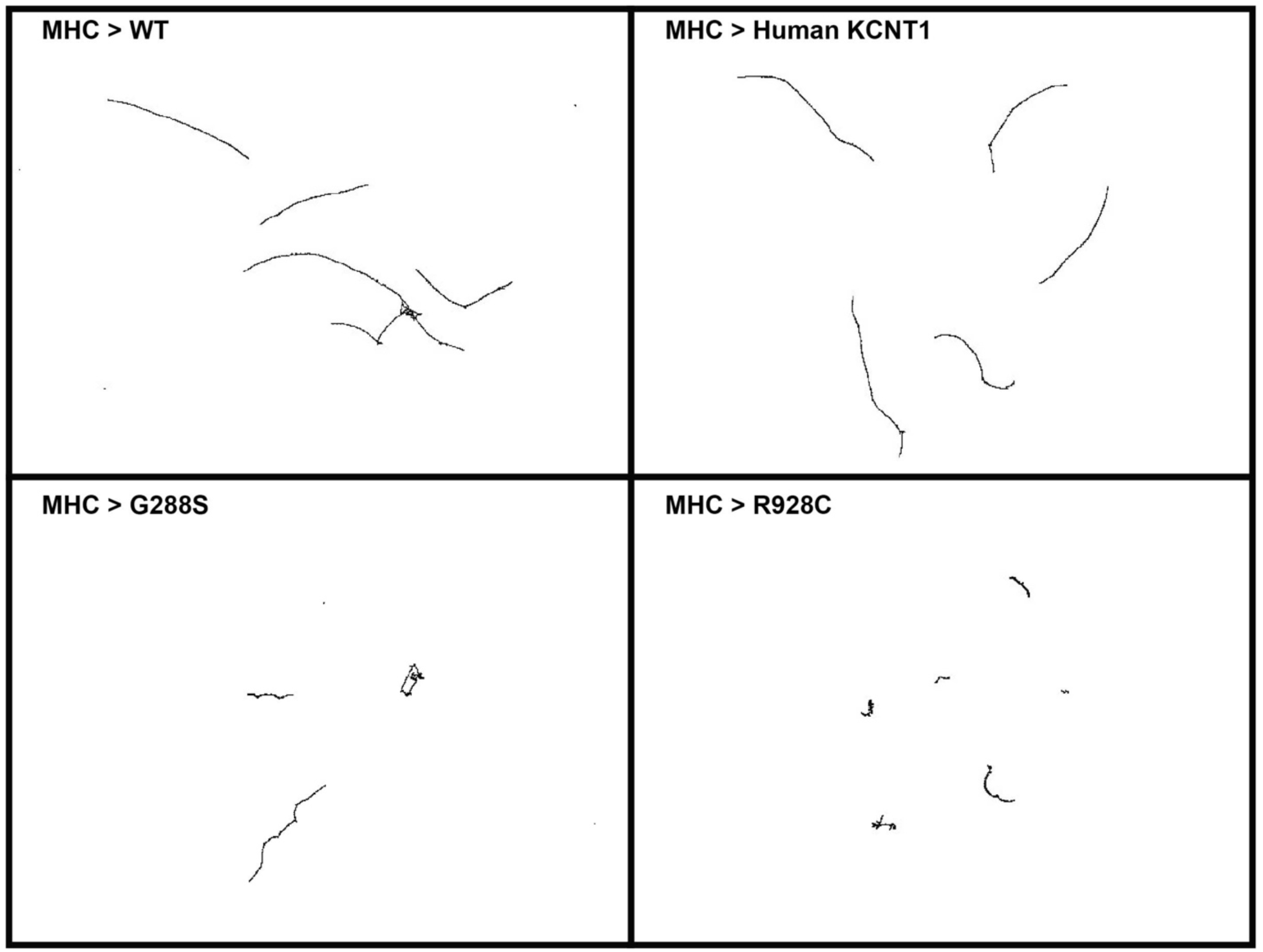
Representative larvae crawling tracks. Larvae expressing mutant hKCNT1 channels in muscles show significant reduction in crawling distance compared to controls.

**Supplementary Figure 3:**
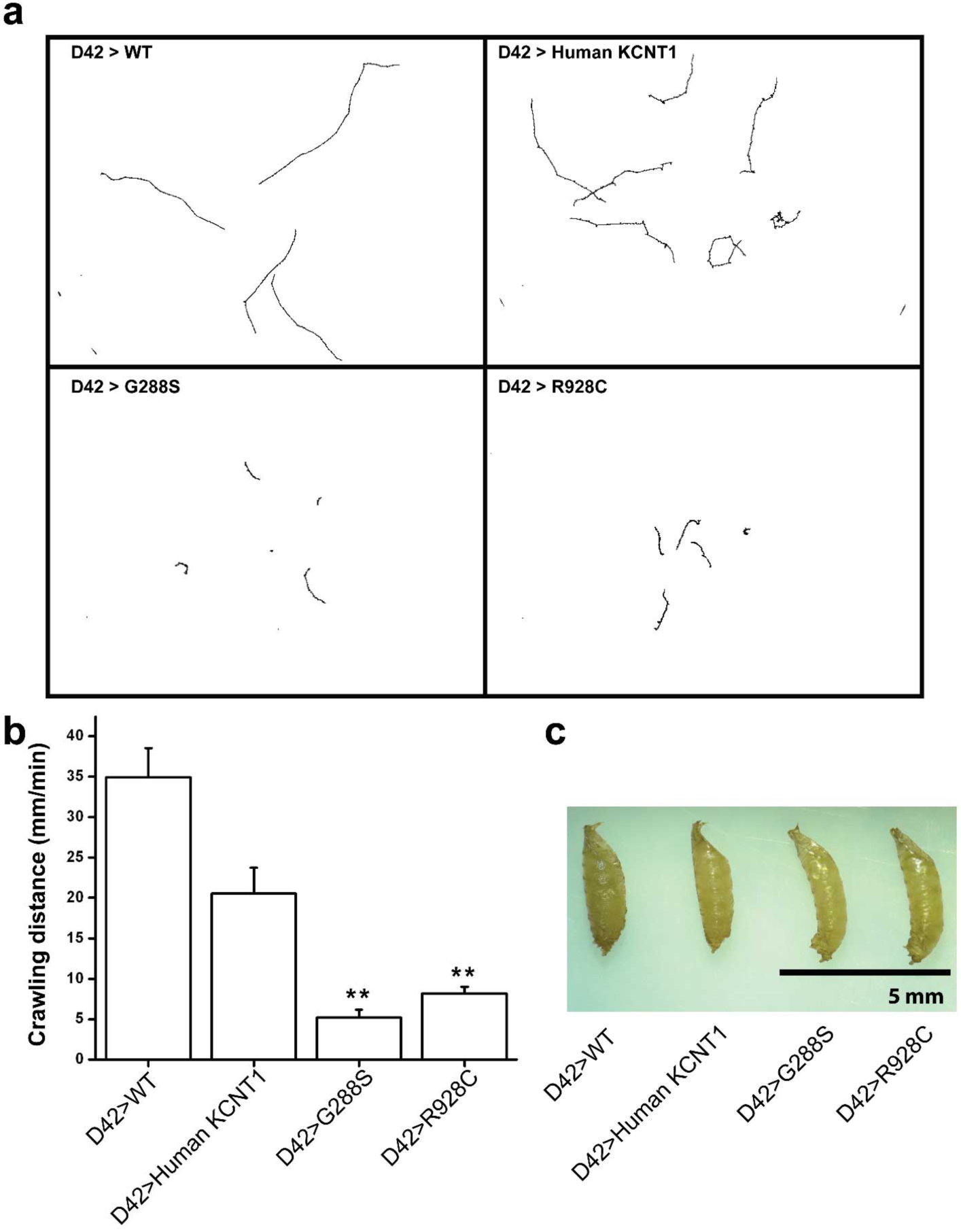
Motor neuronal expression of mutant hKCNT1 channels alter pupal and adult wings morphology and reduces third-instar larval crawling activity. (**a**) Representative larvae crawling tracks. (**b**). (**c**) Motor neuron expression of mutant hKCNT1 channels significantly reduce average crawling distance compare to controls. (**c**) Representative images of pupal morphology. Larvae expressing mutant hKCNT1 channels show bent pupa shape. At least twenty larvae were used per genotype (10 males and 10 females). All larvae were reared at 22 °C for 5 days then at 25 C° for 2 days. Error bars show S.E.M. \*\**P* < 0.01 one-way ANOVA, Turkey HSD *post hoc test*.

**Supplementary Figure 4:**
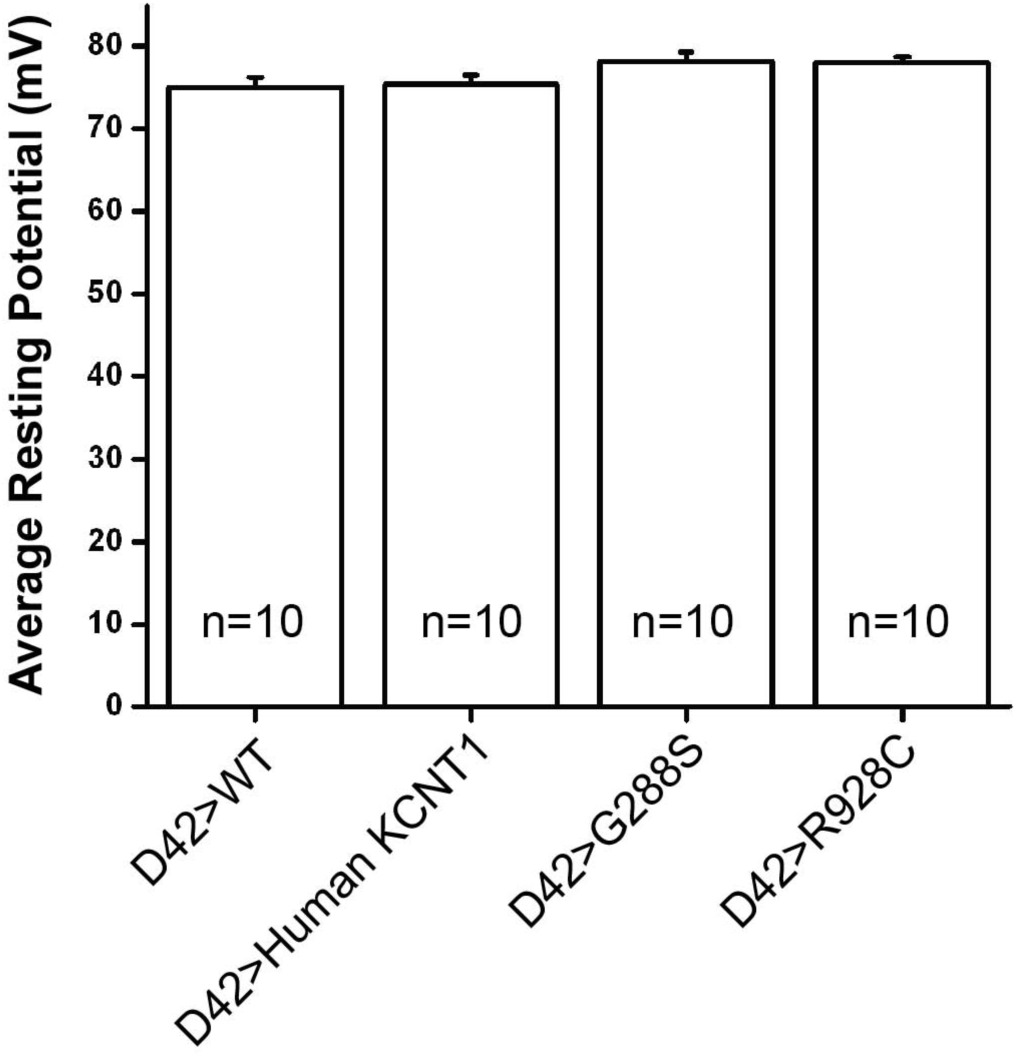
Motoneuronal expression of mutant hKCNT1 channels did not affect membrane resting potential. Both controls and larvae expressing mutant hKCNT1 channels in motoneurons have similar membrane resting potentials.

**Supplementary Figure 5:**
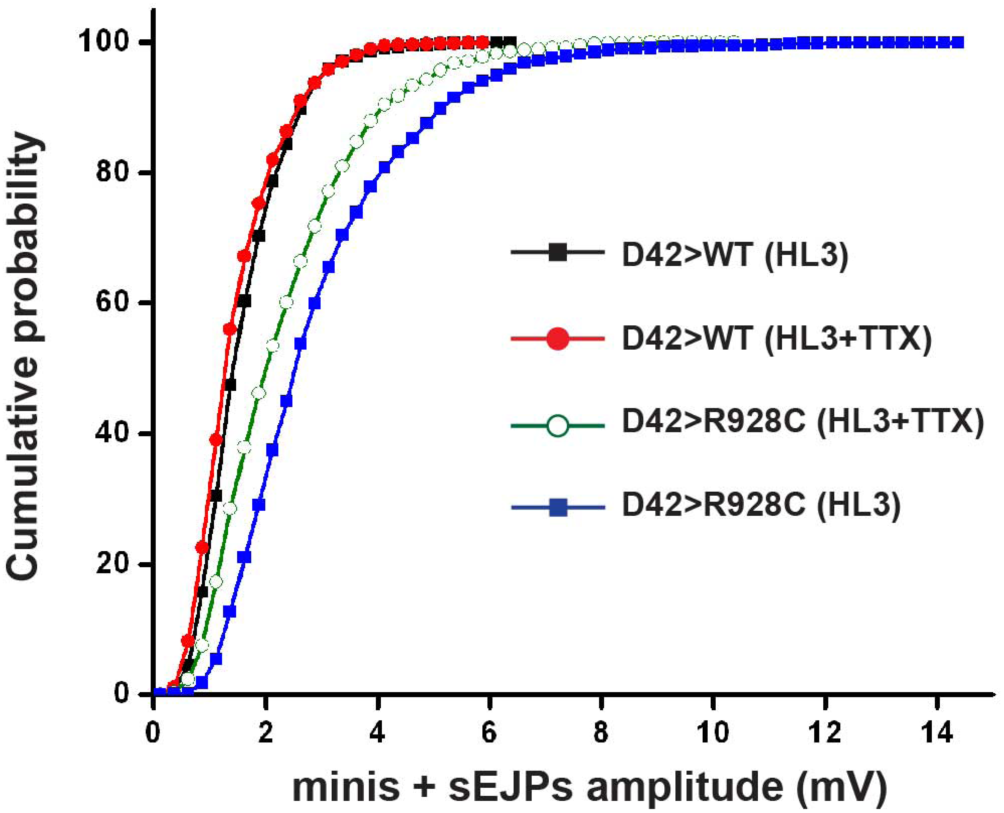
Blocking voltage-gated Na^+^ channels with TTX reduced the number of extremely large sEJPS but did not abolish sEJPs in larvae expressing mutant hKCNT1 channels. Cumulative probability plot of minis plus sEJPs shows that TTX reduced the number of the extremely large sEJPs (>10 mV), but did not eliminate the sEJPs.

**Supplementary Figure 6:**
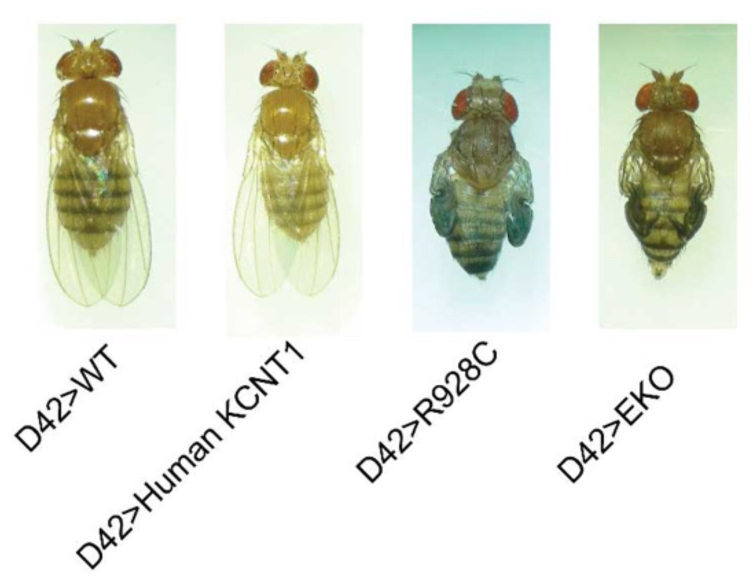
neuronal expression of the mutant KCNT1 Channels causes folded wing phenotype. Representative images of adult fly wing morphology. All larvae were reared at 18 °C until eclosion.

### Supplementary Movie legends

**Movie 1: Muscle expression of mutant hKCNT1 channels hinders larvae crawling activity.** Representative crawling activity assay videos of controls and mutant KCNT1 larvae. Larvae expressing mutant hKCNT1 channels in muscle show significant reduction in crawling activity compared to controls. Twenty larvae (ten males and ten females) were used per genotype, and all larvae were aged at 25 °C prior to testing.

**Movie 2: GABAergic expression of mutant hKCNT1 channels increase the number of seizing flies and seizing duration in adult flies.** Representative bang-sensitivity behavioral assay videos. Flies expressing mutant hKCNT1 channels in inhibitory neurons exhibit seizure phenotypes, such as spinning in the bottom of the vial or flip on their back and remain at that state for one second or more. At least 40 flies were used per genotype, and all larvae were aged at 18 °C until they enclosed then adult flies are aged at 25 °C prior to testing.

**Movie 3: Motoneuronal expression of mutant hKCNT1 channels increase the number of seizing flies and seizing duration in adult flies.** Representative bang-sensitivity behavioral assay videos. Flies expressing mutant hKCNT1 channels in motoneurons exhibit seizure phenotypes, such as spinning in the bottom of the vial or flip on their back and remain at that state for one second or more. 120 flies were used per genotype, and all larvae were aged at 18 °C until they enclosed then adult flies are aged at 25 °C prior to testing.

**Movie 4: Larvae expressing mutant hKCNT1 channels in motoneurons shows uncoordinated contraction of left and right abdominal muscle hemisegments.** Representative crawling activity assay videos of controls and mutant KCNT1 larvae. Larvae expressing mutant hKCNT1 channels in motoneurons show significant reduction in crawling activity compared to controls, due to unsynchronized muscle contractions. Twenty larvae (ten males and ten females) were used per genotype, and all larvae were aged at 22 °C for 5 days and at 25 °C for two days prior to testing.

**Movie 5: Ca^2+^ waves in larvae expressing mutant hKCNT1 channels in motoneurons shows uncoordinated VNC abdominal ganglia firing.** Representative Ca2+ waves assay videos of controls and mutant hKCNT1 larvae. Larvae expressing mutant KCNT1 channels in motoneurons show significant increase in unsynchronized abdominal ganglia firing. Ca^2+^ waves recording was at 2 frames per second and movie is showing 4X the original speed. Four larvae were used per genotype, and all larvae were aged at 22 °C for 5 days and at 25 °C for two days prior to testing.

## Methods

### Fly Strains

Fly cultures kept at 12-h light/dark cycle on standard cornmeal food. The following *Drosophila* strains were used: *D42-GAL4*, *GAD1-GAL4*, *GAL80*^*ts*^, *Mhc-GAL4*, *8622*, and *UAS-GCaMP6S* were obtained from Bloomington Stock Center. *UAS-Kir2.1* was gift from G. Davis, *UAS-EKOX2*, gift from B. White, and UAS-KCNT1 transgenic lines, gift from L. Dibbens.

### Larvae crawling behavior assay and analysis

Wandering third-instar larvae were washed with double distilled water and placed on 150mm petri dish containing 1% agarose. The larvae locomotion behavior was recorded for 1 min at 30 frames/s by Fujinon DF6HA-1B camera and FlyCapture software. Videos were analyzed using ImagJ and wrMTrck plugin as previously descried^59^.

### Calcium imaging

Third-instar larvae were dissected and ventral nerve cord, which contains motoneurons, was viewed using Olympus BX61 compound microscope with 10X air lens. Calcium waves were recorded using ORCA-R^2^ CCD camera (Hamamatsu) and CellSens Dimension 1.9 software. ImageJ software was used for calcium waves analysis.

### Electrophysiology

Recordings were made from muscle 6 in abdominal segments 3 and 4 from third-instar larvae as previously discribed^35,36^. Recordings were made with a modified HL3 saline^60^ containing 70mM NaCl, 5mM KCl, 10mM MgCl_2_, 10mM NaHCO_3_, 5mM trehalose, 115mM sucrose, 5mM HEPES, and 1.2mM CaCl_2_. For true mini was recorded with HL3 solution plus 1 µM TTX and 10 µM CdCl_2_. Recording microelectrodes were fabricated using P2000g puller (Sutter instruments) and filled with 3M KCl. Signals were recorded by Axoclamp B2 amplifier and pCLAMP 6 software (Molecular devices). Severed segmental nerve was sucked by stimulation microelectrode and stimulated by Master-8 pulse generator and iso-flex stimulus isolator (A.M.P.I.).

### Fly bang-sensitive assay

Experiment was done as previously descried with some modification^29,30^. Briefly, flies were collected immediately after eclosion, kept at 25 °C, and mechanically stimulated with mini vortexer (VWR Scientific Products) at power setting 10. Flies behavior was recorded by SONY (HDR-SR11) camcorder. Flies that flipped on their back and remained at that position for at least one second were considered to be exhibiting seizure activity.

### Immunocytochemistry

Wandering third-instar larvae were dissected in HL3 solution and fixed for 2h in 4%paraformaldehyde. Fixed larvae were washed 3X 15 min each with PBST (0.2 TritonX-100). Larvae were incubated overnight at 4 °C with primary antibody with the appropriate dilution. The following antibodies were used: guinea pig anti-Shaker 1:500 (gift from Kyunghee Koh), rabbit HRP-488 1:500. Alexa-conjugated secondary antibodies (1:500) were used. Images were acquired with Leica SP2 confocal microscope using a 63X oil immersion objective.

### Statistical analysis

Data were analyzed as described in the figure legends using Origin software version 6.0.

## Acknowledgements

We thank an internal fund for supporting this line of research from the University of Missouri and Division of Biological Sciences and a postdoctoral fellowship from King Abdullah International Medical Research Center, Riyadh, Saudi Arabia for the support of SNE. We thank Kyunghee Koh for the gift of the Shaker antibody, Leanne Dibbens for sharing fly lines, Weijie Liu for her initial contributions to this project, and the members of the Zhang lab for helpful discussions.

## Author contributions

SNE, GTD, BZ designed the experiments; DD helped design the molecular cloning of KCNT1 channels; SNE, GTD, PS, and BZ collected and analyzed data; SNE and BZ wrote the manuscript, GTD, SP, and DD revised or made comments on the manuscript.

## Competing financial interests

This work described her has been filed for a provisional patent application.

